# Efficient simulation of 3D reaction-diffusion in models of neurons and networks

**DOI:** 10.1101/2022.01.01.474683

**Authors:** Robert A. McDougal, Cameron Conte, Lia Eggleston, Adam J.H. Newton, Hana Galijasevic

**Affiliations:** Department of Biostatistics, Yale School of Public Health, New Haven, CT; Center for Medical Informatics, Yale University, New Haven CT; Department of Neuroscience, Yale School of Medicine, New Haven, CT; Department of Statistics, The Ohio State University, Columbus OH; Yale College, Yale University, New Haven CT

**Keywords:** reaction-diffusion, computer simulation, 3D, multi-scale modeling, reusability

## Abstract

Neuronal activity is the result of both the electrophysiology and chemophysiology. A neuron can be well represented for the purposes of electrophysiological simulation as a tree composed of connected cylinders. This representation is also apt for 1D simulations of their chemophysiology, provided the spatial scale is larger than the diameter of the cylinders and there is radial symmetry. Higher dimensional simulation is necessary to accurately capture the dynamics when these criteria are not met, such as with wave curvature, spines, or diffusion near the soma.

We have developed a solution to enable efficient finite volume method simulation of reaction-diffusion kinetics in intracellular 3D regions in neuron and network models and provide an implementation within the NEURON simulator. An accelerated version of the CTNG 3D reconstruction algorithm transforms morphologies suitable for ion-channel based simulations into consistent 3D voxelized regions. Kinetics are then solved using a parallel algorithm based on Douglas-Gunn that handles the irregular 3D geometry of a neuron; these kinetics are coupled to NEURON’s 1D mechanisms for ion channels, synapses, etc. The 3D domain may cover the entire cell or selected regions of interest. Simulations with dendritic spines and of the soma reveal details of dynamics that would be missed in a pure 1D simulation. We describe and validate the methods and discuss their performance.

## Introduction

The brain’s behavior in health and disease is most naturally observed at the level of functional outcomes, but these outcomes are often indirect consequences of subcellular chemical kinetics (e.g. oxygen and ATP in stroke; amyloid beta and tau in Alzheimer’s Disease). The connection between these two scales is non-intuitive due to the many nonlinear-interactions within the brain (e.g. action potentials, networks). Dedicated tools like MCell (Stiles et al., 1998) and STEPS (Hepburn et al., 2012) enable highly-detailed 3D simulation of parts of neurons to entire cells, enabling the study of microdomains (see e.g. Basak and Narayanan (2018)) and other highly localized phenomena but with limited ability to extend to the full cell or a network of neurons to study the implications of these localized dyanmics on a broader scale.

The NEURON simulator (Hines et al., 2019) has long supported simultaneous simulation of chemical dynamics and networks of neurons, originally through re-purposing MOD files – traditionally used for ion channel kinetics – and later through the introduction of a dedicated Python-based reaction-diffusion specification (McDougal et al., 2013b). These early methods were most applicable to phenomena that behave analogously to electrical signaling, such as when a wave of elevated calcium concentration spreads over a large region of the dendritic tree (e.g. Neymotin et al. (2015)). Even for these large scale phenomena, the 1D approximation breaks down in regions where the cell is not radially symmetric (e.g. the predicted curvature of a calcium wave front where the dendrite meets the soma as in Fig. 1A) and is inappropriate for smaller scale phenomena on the same spatial scale as the dendrite diameter (e.g. diffusion between neighboring spines as in Fig. 1B).

**Fig. 1.**
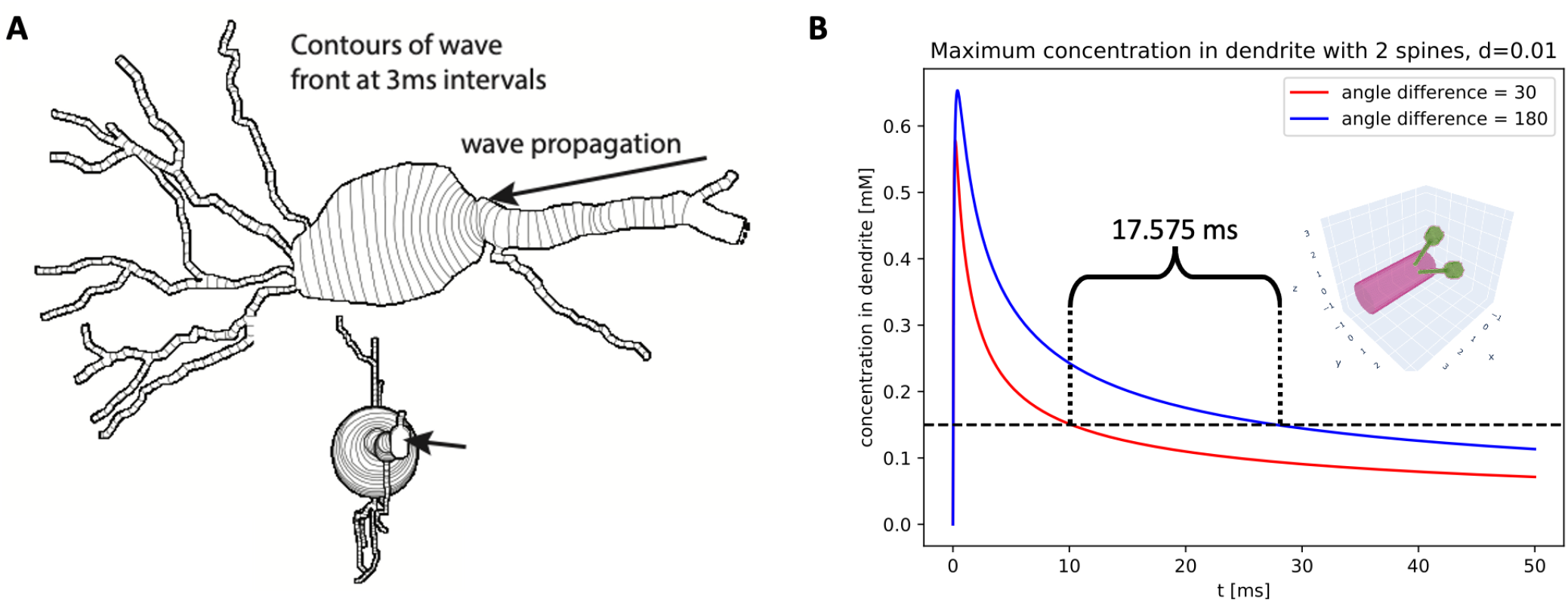
Chemical concentration within neurons is affected by the 3D shape of the cell. (A) An intracellular wave entering the soma may curve, slow, and in extreme cases fail to propagate. (B) Diffusion of ligand from neighboring spines (red; illustrated as inset) leads to different trajectories of higher peak dendrite concentrations than from spines opposite each other on the dendrite.

We have developed a set of approaches with implementations freely available in the development version of NEURON – to efficiently address the need to incorporate 3D intracellular dynamics for subcellular compartments, whole cell and network models. Combining local discretizations and preserving segment mappings accelerates the Constructive Tessellated Neuronal Geometry algorithm (CTNG; McDougal et al. (2013a)) for generating a 3D volume consistent with a neuron point-and-diameter 3D reconstruction, of the sort available via e.g. NeuroMorpho.Org (Ascoli et al., 2007). Reaction-diffusion (rxd) kinetics are specified as for 1D simulations, with selected regions of interest simulated selected for 3D simulation via a single line function call, while other parts simulated in 1D. The 3D regions of interest are voxelized and any overlapping 3D regions are connected together and with neighboring 1D regions. Threaded, deterministic simulation is enabled using an irregular boundary extension of Newton et al. (2018)’s operator-splitting parallelized Douglas-Gunn Alternating Direction Implicit method (Douglas and Gunn, 1964). Ion channel activity is based on concentrations at the surface of the cell, and ions enter the cell through the surface voxels. Single cell results are validated by comparison to analytic solutions, by comparison of 3D results with other tools, and by comparison of hybrid 1D-3D simulations with pure 3D simulations.

## Methods

Methods and results are described for a development version of NEURON 8.1, although the initial version of most of these methods was introduced in NEURON 7.7. The source code is available at github.com/neuronsimulator/nrn, installers for major platforms are available at neuron.yale.edu, and NEURON can also be installed for linux and macOS via `pip install neuron`. The voxelization algorithm is written in a mix of Python, Cython, and C/C++. The interface code is written in Python. For performance reasons, all NEURON reaction-diffusion code used during an active simulation is written in C/C++.

For analyses requiring many simulations, simulation and visualization were split into separate scripts with each simulation’s data stored in a SQLite database. To be robust against the possibility of interrupted calculation, simulation scripts checked the database to see if a given set of parameters had already been tested before running the simulation. Graphs were rendered using plotly (for 3D images), plot-nine/ggplot, and matplotlib.

Python code for all figures in this manuscript is available on ModelDB (McDougal et al., 2017) at modeldb.yale.edu/267018.

### Voxelization

3D simulation requires the specification of a 3D domain, typically defined by a mesh (e.g. in VCell) or a boundary (e.g. MCell, Smoldyn). Neuron morphologies, by contrast, are typically reconstructed using a series of (*x, y, z; d*) optical measurements with tree-structured connectivity rooted at the soma, which is sometimes a special case with an outline, typically in 2D. (A neuron’s morphology is a computer science tree in the sense that every non-root section has exactly one parent section, namely the connecting section that is closer to the root.) This information is sufficient for electrophysiology simulation where the space constant is typically on the order of tens of microns, but under-determines the 3D structure for chemical simulation. Several algorithms have been proposed to generate consistent geometries, including our Constructive Tessellated Neuronal Geometries (CTNG) algorithm (McDougal et al., 2013a) and others (e.g. Lasserre et al. (2011), Mörschel et al. (2017)). The full CTNG method is described in our previous paper, but in brief consecutive point-diameter measurements are interpreted as defining the frustum of a right circular cone. Neighboring frusta are joined using clip spheres, with a clipping rule that depends on the taper of frustra and the angle of intersection. Soma outlines are approximated using sheared frusta with dendrites attached to the soma extended to the soma axis to avoid any gaps from the assumption of local radial symmetry given a 2D soma outline. NEURON’s Import3D tool stores soma outline points in a Python dictionary; these soma outlines are not used in pure electrophysiology simulations, but the voxelization algorithm checks each section against the dictionary to see if it should be treated as a sequence of frusta or if there is a 2D outline to use.

To accelerate CTNG voxelization and to facilitate its use in simulations incorporating one-dimensional electrophysiology dynamics, we enhanced the original implementation in several ways: (1) additional interpolated points are inserted at electrophysiological compartment (“segment” in NEURON) boundaries so every frusta belongs to exactly one compartment; (2) each electrical compartment is voxelized separately, thus preserving the relationship between voxels and electrical compartments; (3) each frusta and joining sphere is voxelized separately, exploiting convexity to rapidly identify all the relevant voxels; and (4) voxelized meshes are merged together, with voxels being assigned to the segment closest to the root of the electrophysiological tree (typically the soma).

In CTNG, the constructed 3D volume of the neuron is the union of frusta and spheres clipped by planes. These component objects are voxelized using a modified flood-fill algorithm, starting from the center of an end-face for a frustum or the center of the sphere. In the case of a clipped sphere with a small angle, the resulting wedge may be very small, causing all corners of the voxel to be outside. In this case, additional points in the sphere are tested until a voxel is found with at least one interior corner; this voxel is then used as the seed for the flood-fill. Since the spheres serve to smooth the joins between neighboring frusta, in practice they comprise a small percentage of the total voxels, so this search introduces minimal overhead.

The flood fill is propagated through the surface of the shape as much as possible, using the convexity of the objects to automatically fill in interior voxels between two surface points. This is done by traversing rows of voxels perpendicular to the *y, z* plane. No matter the orientation of the object, any row of voxels that contains part of the object contains one or more surface voxels at each endpoint of the row’s intersection with the object. The modified flood fill calculates these endpoints along with any additional surface voxels in the row, and then fills in any non-surface voxels between the endpoints as interior voxels. To find all the rows intersecting the object, the flood fill searches all the rows bordering an intersecting row, using the endpoints of the original row as “guesses” to retain information about the surface and expedite finding endpoints for the surrounding rows. The signed distance to each surface voxel corner is also computed and stored throughout the flood fill; as in the original CTNG implementation, these signed distances are supplied to the marching cubes algorithm (Lorensen and Cline, 1987) to approximate the surface with a triangular mesh.

To approximate the surface, the marching cubes algorithm requires that at least one corner of a voxel containing the surface be outside the object and at least one corner be inside. NEURON generates a warning suggesting a smaller *dx* (the length of a voxel edge) if any frustum length or diameter is less than the largest distance that fits within a voxel 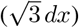. Some publicly available reconstructions are sampled with very little distance between the 3D points, leading to a very small suggested value of *dx* and as noted in McDougal et al. (2013a), sometimes a bumpy 3D reconstruction; in this case, subsampling the 3D points before loading the morphology into NEURON avoids both the bumpiness and the recommendation of a small *dx*. The extra segment boundaries added by a very high (per unit length) value of `nseg` (the number of electrical compartments in a Section) can produce a similar effect; in this case, the solution is to reduce `nseg` to a value appropriate for the electrical space constant. In NEURON, an appropriate choice of `nseg` can be determined for each Section based on the so-called `d_lambda` rule (Hines and Carnevale, 2001).

The areas of the triangles in the surface mesh are summed to estimate surface area, and the portion of each surface voxel inside the object is estimated to be the fraction of test points inside the object. As the voxel has been identified as a surface voxel, at least one corner is inside, and thus the volume estimate will never be 0. Test points are sampled on a uniform grid in 1 + `options.ics_partial_volume_resolution` steps in each direction along the voxels edge, starting and ending at a voxel corner. NEURON versions 7.7 - 8.0 used an alternative rule for estimating partial volumes using dynamic subsampling, however the approach described above and used beginning in NEURON 8.1 is simpler and provides better scaling.

As neurons occupy a small fraction of the volume of their bounding box (1.498% ± 3.406% for the neurons in the Random Realistic Neuron Morphologies Section voxels are stored as a set of locations (*i, j, k*) within an imaginary grid comprising a padded bounding box. Thus, memory usage to store the discretization is proportional to the volume of the neuron not to the volume of the bounding box. Likewise, NEURON’s simulation times scale proportionally to the number of voxels in the cell, not the number of voxels in the bounding box.

Discretization into a 3D grid happens as needed, allowing interactive changes to grid parameters, morphology, etc without the overhead of re-voxelizing the cell. The mesh is typically generated on the first request for a pointer (e.g. for recording concentration at a point), or when the simulation is initialized. NEURON’s internal counters for structure or diameter changes are monitored for subsequent changes at each initialization, pointer request, or simulation step, and the morphology is re-discretized if needed; such re-discretization is expected to be rare in practice as NEURON models typically assume cells do not change shape or size during simulation.

### Model Specification

NEURON’s basic reaction-diffusion model specification, introduced in McDougal et al. (2013b), is independent of numerical simulation details such as whether the model is to be simulated deterministically or stochastically or in 1D or 3D. Readers are directed to the 2013 paper or for a more complete and updated treatment to the relevant section of the online NEURON documentation (nrn.readthedocs.io/en/latest/rxd-tutorials) for full details, but in brief: domains of a cell are specified in Python using `rxd.Region`, chemical species and their properties using `rxd.Species`, and chemical reactions using `rxd.Reaction`, `rxd.Rate`, or `rxd.MultiCompartmentReaction`. The classes `rxd.Parameter` and `rxd.State` allow fixed values that change with location and non-diffusing state variables, respectively. Dynamics at a specific point (e.g. localized pump) are specified using `node.include_flux` where `node` represents the spatial compartment and the flux is measured in changes in *mass*. Using mass changes instead of concentration changes allows the same amount of a substance to enter the cell regardless of the spatial discretization. To specify that all reaction-diffusion kinetics should be simulated in 3D, call `rxd.set_solve_type(dimension=3)`.

NEURON automatically translates the Python kinetics specification into C and compiles them for use during simulation as described in Newton et al. (2018).

### 3D Simulation

Intracellular 3D regions are simulated using a generalization of the parallel Douglas-Gunn based method for 3D extracellular simulation of Newton et al. (2018). Unlike the extracellular case which is simulated coarsely enough that the morphological details can be subsumed into an effective volume fraction, in intracellular simulation the voxels are necessarily much smaller and need to respect the 3D boundary of the cell. As described in the Voxelization Section, voxels may have widely varying amounts of surface, and each voxel must be associated with a specific electrical compartment (with a “segment” in NEURON terminology).

The conceptual algorithm for integration is the same as in Newton et al. (2018), however with voxels only existing inside the cell membrane, the number of voxels in any row is no longer necessarily the same. This variation means that although the memory locations for concentration in a particular voxel and in the voxel above it are fixed for a simulation, the offset between voxels and the voxels above them varies throughout the cell and cannot be calculated using a simple arithmetic expression. To work with this irregularity, the indices of every voxel in each line are precomputed at initialization. To find neighbors, NEURON constructs a dictionary (hash array) keyed by the (*i, j, k*) voxel location with the value of the index of the voxel in memory. Lines in each direction are formed by starting from an arbitrary voxel, backtracking to the beginning of the line (e.g. if forming the lines parallel to the *x* axis, we successively check for the presence of (*i* – 1, *j, k*), (*i* – 2, *j, k*), … until such a voxel is not in the dictionary), and then recording the indices of the voxels in the line until there is no next voxel indexed in the dictionary. Although Python is used to calculate the indices comprising each line, the results are cached and transferred to C++ code that uses them during integration. This process is repeated if and only if the 3D structure is changed (e.g. more segments, different diameters, …).

All memory indices are relative to a given `rxd.Species`, (or `rxd.Parameter` or `rxd.State`), each of which has its memory allocated independently, allowing for one to be added or removed without requiring memory for the others to be reallocated.

Fixed step integration proceeds using the two-phase operator-splitting approximation as in (McDougal et al., 2013b): reactions and fluxes are calculated using an implicit method first and then diffusion is calculated with DG-ADI, also an implicit method. This introduces a source of error that converges to 0 as *dt* → 0, and has the advantage of keeping the matrices that need to be inverted (a 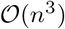 task in the general case) small, involving only one location’s concentrations for each reaction matrix or one line for each diffusion matrix. No calculations are done for memory associated with `rxd.Parameter` objects, only reactions are calculated for `rxd.State` objects, and both reactions and diffusions are calculated for `rxd.Species` objects. Fluxes from ion channels specified with MOD files are converted into mass changes per segment and then distributed proportionally across the surface voxels assigned to the segment by voxel surface area. Here, it is assumed that every segment has surface voxels. All diffusion calculations explicitly incorporate the effect of voxel by voxel interior volume as voxels with surface do not have their entire volume inside the cell.

Variable step integration uses the CVODE solver from the SUNDIALS suite (Hindmarsh et al., 2005) with all NEURON rates of change (membrane potentials, ion channel states, reactions and diffusion) represented in one derivative vector. The approximate Jacobian used for the reaction-diffusion part of the problem is a permutation of a blockdiagonal matrix, where each block includes the full reaction Jacobian for a given spatial location but only the diagonal part of the diffusion Jacobian. This simplifying approximation allows the approximate Jacobian to be quickly invertible at a tradeoff of a decrease in accuracy, potentially forcing smaller timesteps than CVODE might use with the exact Jacobian.

NEURON concentrations are tied to segments and the surface nodes are assigned the concentration. In general, there are multiple surface nodes per segment. To address this, the segment concentrations are updated at each time step with the weighted average concentration from the segment’s surface voxels. In some cases, using only the surface nodes can cause an artificially high concentrations due to relatively few 3D voxels diffusing with a single 1D segment. This effect can be mitigated by using all 3D nodes to calculate concentrations with `options.concentration_nodes_3d = “all”`.

Multiple threads, specified with `rxd.nthread(n)` where n is the number of threads, may be used to accelerate intracellular simulation. Load balancing is achieved using a longest-processing-time first greedy algorithm based on the line length for each direction, which is guaranteed to have no worse than 4/3 the optimal time (Graham, 1969).

### Hybrid simulation

For performance reasons and to better support simulations with narrow dendrites that would otherwise require a small *dx*, a Python iterable of Sections (list, set, …) may be provided when specifying the simulation dimension to indicate which Sections are being set, e.g. `rxd.set_solve_type(apicals, dimension=3)`. Chemical simulations within a given Section must either be in 1D or 3D (i.e. this cannot vary by Region), but each Section can be set independently. Capturing the 3D nature of the dynamics within a Section generally requires that its neighboring Sections also be in 3D as otherwise all the incoming diffusive fluxes from neighboring Sections are the same regardless of 3D location. As with specification of the full model, multiple dimension specifications for a given Section are allowed, with the last one taking effect.

#### Boundary identification

At initialization, each `rxd.Region` identifies which of the sections it contains are to be simulated in 1D and which are to be simulated in 3D. If a region does not contain both sections to be simulated in 1D and in 3D, no additional analysis is done and the simulation is purely in the specified dimension. When both dimensions are present, for every given `rxd.Species` instance, every 3D section’s parent, if it exists, is checked to see if it is on the 1D section list. Likewise, every 1D section’s parent is checked to see if it is on the list of 3D sections. In recommended usage each cell has its own `rxd.Species` instance for a given conceptual molecule (e.g. each cell would have a `self.ca` for its internal calcium concentrations), the search space is constrained to a given cell. Within any given `rxd.Region` containing both 1D and 3D sections, there may be zero (if the 1D and 3D sections are not contiguous), one, or arbitrarily many places where 1D and 3D sections meet.

For each case where 3D and 1D sections meet, boundary voxels are identified by finding the voxels belonging to the 3D section and its spherical endcap that intersect the plane perpendicular to the line segment defined by the two (*x, y, z*; *d*) points at the appropriate edge of the 1D section. In particular, this algorithm requires that the boundary or boundaries must occur at either end of a NEURON Section, not in the middle. Each 1D-3D juncture potentially has many boundary voxels, depending on the 3D discretization. Boundary voxel identifiers and distances to the 1D boundary nodes are computed at initialization and cached in a data structure passed to the C++ compute engine via `ctypes`. Any subsequent changes to the morphology trigger recalculation of the discretization – including identification of boundary voxels – at the next initialization, advance, or node request event.

#### Simulation

At the beginning of each timestep, fluxes between 1D and 3D boundary compartments are computed according to the finite volume method and Fick’s laws: fluxes are proportional to the concentration gradient and inversely proportional to the 1D distance between the centers of the compartments. The 1D and 3D regions are then advanced independently, applying the fluxes as appropriate, thereby weakly coupling them. This weak coupling introduces minimal performance overhead, but at the cost of reduced numerical stability, thereby potentially requiring a smaller timestep (see e.g. Benedikt and Drenth (2019).)

### Random realistic neuron morphologies

To assess performance on realistic morphologies, we identified 21 random reconstructions from NeuroMorpho.Org (Ascoli et al., 2007) with metadata indicating realistic diameters and a 3D reconstruction. These were obtained by querying NeuroMorpho.Org’s “Browse by Random” tool, once for 50 random cells and once for 10 random cells, and filtering for those meeting the stated criteria. The randomly selected morphologies as identified by their NeuroMorpho.Org name are: `9CL-IVxAnk2-IR_ddaC` (Nanda et al., 2018), `29-1-8` (Martinez-Canabal et al., 2013), `64-8-L-B-JB` (Ehlinger et al., 2017), `243-3-39-AW` (Nguyen et al., 2020), `2017-25-04-slice-2-cell-2-rotated` (Scala et al., 2019), `070601-exp1-zB` (Groh et al., 2010), `160524_7_4` (Kunst et al., 2019), `15892037` (Takagi et al., 2017), `AE5_EEA_Outer-thirds_DG-Mol_sec1-cel4-aev5me` (de Oliveira et al., 2020), `AM61-2-1` and `AM81-2-3` (Trevelyan et al., 2006), `B4-CA1-L-D63×1zACR3_1` (Canchi et al., 2017), `Dnmt3bKO-cell-8 and WT-iPS-derived-cell-12MR` (Tarusawa et al., 2016), Fig5C (Herget et al., 2017), `glia_4090` (Helmstaedter et al., 2013), `KC-s-4505762` (Takemura et al., 2017), `Mouse_CA2_Ma_Cell_5` (Helton et al., 2019), `RatS1-6-107` (Nogueira-Campos et al., 2012), `RP4_scaled` (Weiss et al., 2020), and `WT-mPFC-A-20X-3-2` (Juan et al., 2014).

### Timings

All reported times are based on measurements on Yale’s Farnam HPC’s general partition, which has a mix of mostly Intel Xeon E5-2660 v3 CPUs with 119 GiB memory per node and some Xeon 6240 CPUs with 181 GiB memory per node.

## Results

### Validation

#### Convergence on a cylinder and the role of voxel refinement

To assess convergence of surface area and volume calculations, we began by considering a cylinder of diameter 2 μm and length 5 μm. Cylinders, unlike neuron morphologies from reconstructions, have analytically known values for surface area and volume; in particular, here the volume is 5*π* μm^3^ and the surface area is 12*π* μm^2^. Errors were measured for one thousand random orientations (specified as (*ϕ,θ*) in spherical coordinates) at *dx* = 2^−1^, 2^−1.5^, 2^−2^, 2^−2.5^, 2^−3^, 2^−3.5^, and 2^−4^ μm without using partial volume resolution on the surface voxels. For reasons that are explored later, we did not choose this to be NEURON’s default behavior; using the defaults produces volume errors typically around 4x lower.

Volume estimates converged with error scaling at approximately 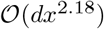, with the average absolute error at *dx* = 2^−1^ being approximately 0.3976 ± 0.4396 μm^3^ and the error at *dx* = 2^−4^ being approximately 4.271 × 10^−3^ ± 8.590 × 10^−3^ μm^3^, a 93.10-fold reduction. This convergence compares favorably to the simpler volume estimating approach of counting the full volume of every voxel that is included in the geometry, which converges at approximately 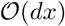. In contrast to that approach, which by definition gives overestimates of volumes, our algorithm gave more underestimates than overestimates (582 out of 1000 test orientations) when dx=0.25 μm, with an average signed error of −4.205 × 10^−2^ μm^3^.

Surface area converged at approximately 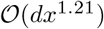: absolute errors at *dx* = 0.5 μm were 2.963 ± 0.313 μm^2^, and at *dx* = 2^−4^ μm were 0.2384 ± 0.0251 μm^2^, an approximately 12.43-fold reduction. For a convex shape such as a cylinder, surface areas estimated using marching cubes are expected to be under-estimates (and this holds for 999 out of 1000 of our test cylinder orientations) as the computed surfaces are planes lying strictly inside the shape; as neurons are not convex, surface areas need not be underestimates in those morphologies.

As interior voxels always contribute their full volume – and therefore do not contribute to volume error – and have no surface, we examined if it was advantageous to use a refined mesh on the surface voxels only to reduce error without incurring the full speed penalty from using a finer global mesh. As explained in the methods, we subdivide surface voxels into VR^3^ subvoxels, compute the volumes and surface areas for each, and sum the results together for the total values for the voxel. Although this approach is similar to increasing the overall mesh resolution, it still depends on the coarser mesh’s determination of which voxels are surface or not, and thus differences may arise in complex morphologies when, for example, small branches pass near each other.

To examine the effect of voxel subdivisions, we held the cylinder orientation constant, parallel to the *x*-axis, and recorded the errors and runtime for various subdivision levels VR. In NEURON, subdivisions used for volume and surface area calculations are independently configurable, using `rxd.options.ics_partial_volume_resolution` and `rxd.ics_partial_surface_resolution`, respectively. In general, as shown in Figure 2, increasing VR provides volume errors and discretization times comparable to using a higher resolution grid but without introducing additional voxels that would significantly increase simulation overhead. Subdividing for surface area calculation did not improve the error for a given discretization time, likely due to this strategy increasing the number of marching cubes to compute since it must process domains that would otherwise be classified as fully interior or fully exterior. As such, NEURON’s default `rxd.options.ics_partial_volume_resolution` of 2 and `rxd.ics_partial_surface_resolution` of 1 are used for all subsequent calculations, i.e. volume calculations use subdivided surface voxels while surface area calculations do not.

**Fig. 2.**
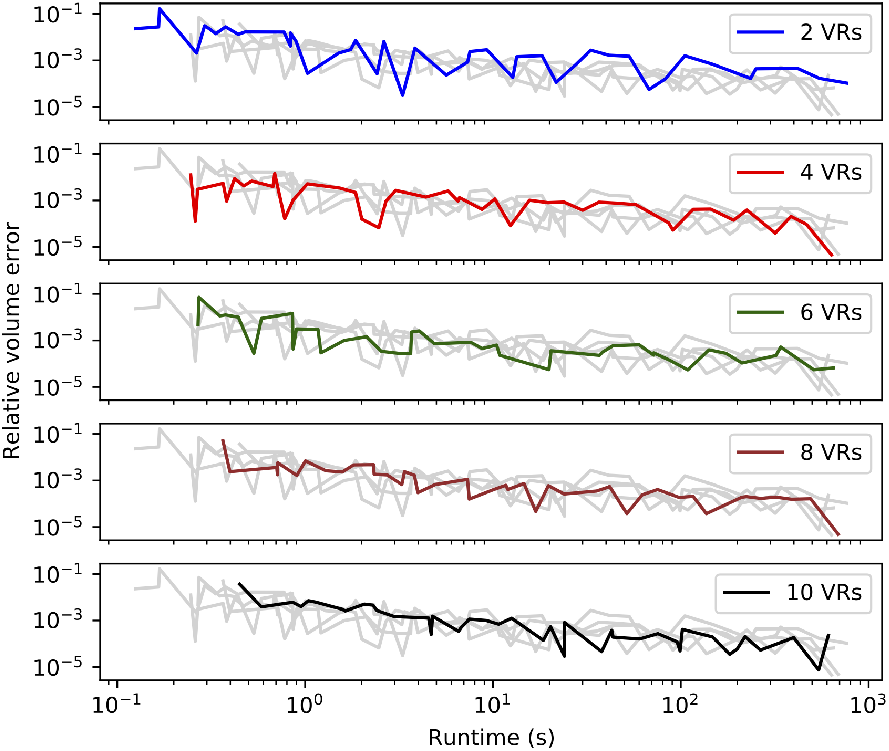
Sub-sampling surface voxels tends to improve the accuracy of the volume estimate for any discretization resolution at the cost of increasing voxelization time. VR is the partial volume resolution, the voxelization times of a cylinder of fixed size and orientation are shown for for 50 different values of dx (from 0.01 μm to 0.5 μm) and 5 values of VR (2, 4, 6, 8, 10). Each subplot highlights one of the VRs, with the others shown in grey. As dx decreases, discretization time increases and relative error tends to decrease, but the error is non-monotonic due to changing alignment of the cylinder with the grids.

#### Convergence of discretization on realistic geometries

To assess the convergence of volume and surface area estimates on realistic morphologies, we used our voxelization algorithm to estimate these values for 21 randomly chosen neuron reconstructions from NeuroMorpho.Org as described in the random realistic neuron morphologies section. For most cells, we tested 12 choices of dx from 0.05 μm to 0.5 μm, omitting the smaller values for large cells that would require prohibitive setup time or memory at those resolutions. As the true surface area and volumes are unknown, we compared each value for a given morphology to the corresponding value calculated with the smallest dx (Figure 3). With dx=0.5 μm, the majority of whole cell morphologies (13 out of 21) had an estimated relative volume error of less than 1% with one having error less than 0.1%. At NEURON’s default resolution of dx=0.25 μm, 15 out of 21 morphologies had a volume error less than 1% with 5 having a volume error less than 0.1%. By dx=0.15 μm, these rates increase to 19 out of 21 and 10 out of 21, respectively. The volume error scaling varies per morphology but scales between 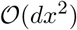 and 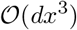 (Figure 3B). With dx=0.15 μm, 10 out of 21 whole cell morphologies had estimated surface area errors less than 1% and two morphologies had surface area errors less than 0.1%. For most morphologies, the surface area error scaled between 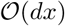 and 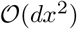 (Figure 3A). Thus, the scaling rates for volume and surface area errors wth realistic neuron morphologies are broadly consistent with the rates observed for the cylinder.

**Fig. 3.**
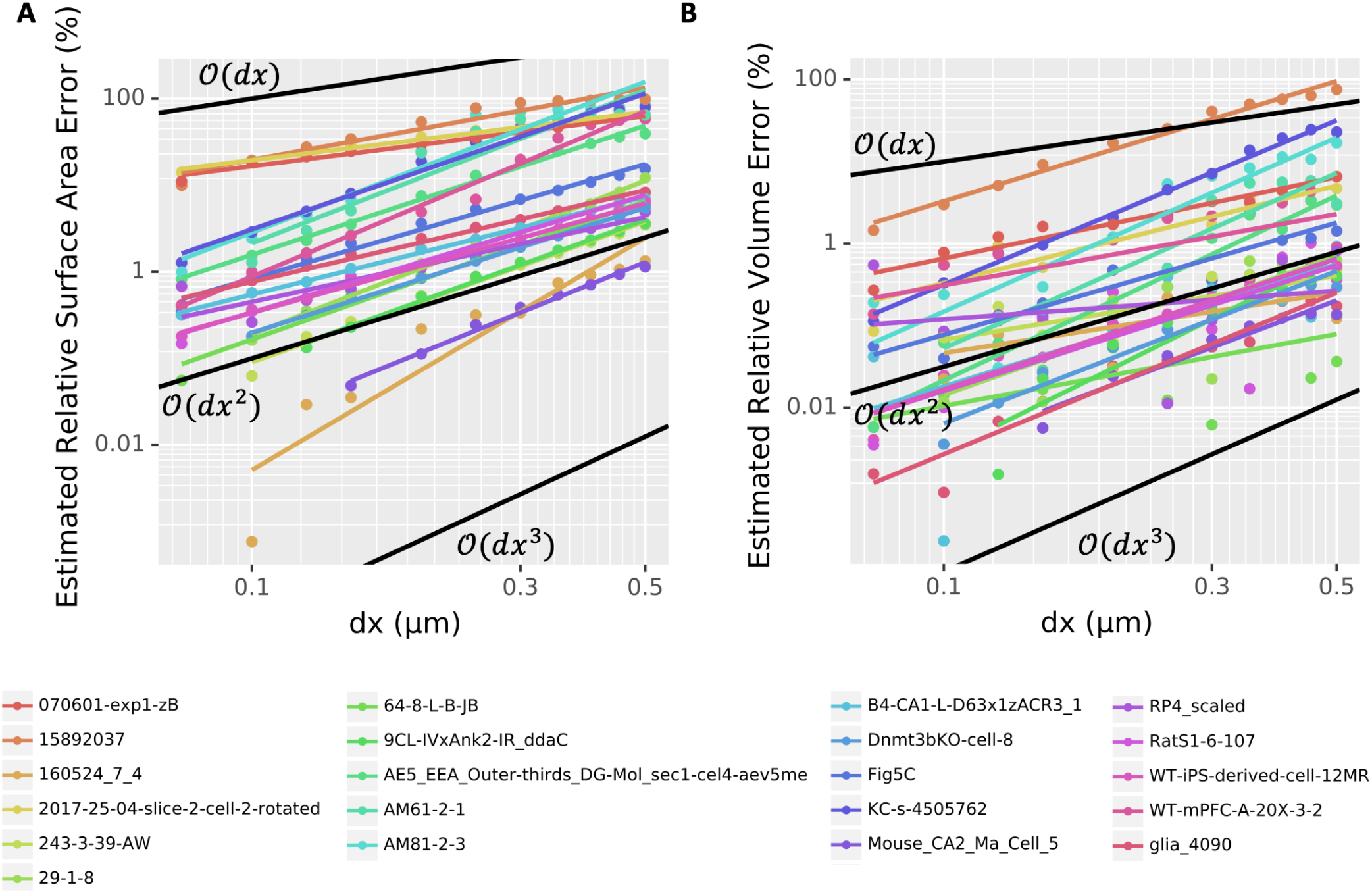
Log-log plots of estimated (A) surface area and (B) volume relative error as a function of dx for the voxelization of 21 entire morphologies (all sections) chosen randomly from NeuroMorpho.Org. Points indicate measured values; colored lines indicate best-fits. Black lines indicate first-, second-, and third-order convergence, as marked, for reference. Note that the *y*-axis scale is different between (A) and (B).

#### Voxel-segment assignment

For currents through the membrane to correctly alter and be modulated by local near-surface concentrations, each voxel must be assigned to the correct segment and surface voxels must be distinguished from interior voxels. To test these classifications, we constructed a simple geometry, consisting of two parallel connected cylinders, with length 5 μm, diameter 5 μm, and 9 segments, and length 5 μm, diameter 1 μm, 5 segments, respectively. We plotted the quarter of the surface-voxels with *x* > 0, *y* > 0, and *z* > 0 in 3D and color-coded by segment, Figure 4A. (The *x* > 0 condition removes the end-face of one of the cylinders.) Visual inspection revealed that our algorithm constructed a continuous surface with no holes and no interior mis-identified voxels. Similar results were found for the other sections of the geometry (not shown), suggesting that the algorithm correctly distinguishes surface and nonsurface voxels. To test the voxel-segment assignment, we projected this image into the *x, y* plane and added markers for the analytically computed segment boundaries (every 5/9 μm for the bigger cylinder and every 1 μm for the smaller cylinder). We additionally added a line segment that passes through the corner of the big cylinder and the midpoint of the first segment of the smaller cylinder which by default in NEURON is assigned a 3D point, and thus this line segment marks the projection of the cone that CTNG adds to join the two cylinders. All segment boundaries aligned with the analytically computed ones and the join cone tapered as expected; Figure 4.

**Fig. 4.**
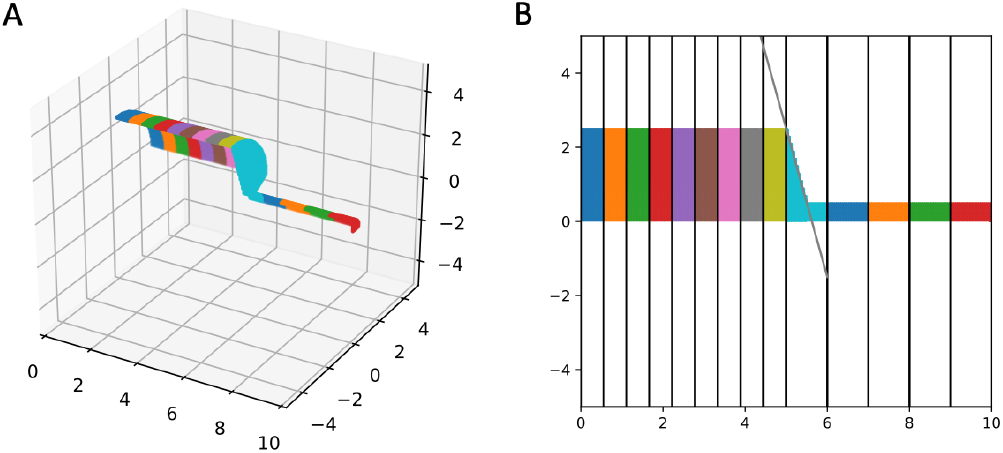
Segment alignment validation. (A) 3D plot and (B) 2D-projection of surface voxels of a morphology with an abrupt change in diameter and a change in segment length, colored by segment. Vertical lines in (B) illustrate the locations of the 1D segment boundaries, which align with the 3D surface nodes. The diagonal black line connects the edge of the last wide segment with the top-middle of the first narrow segment and matches the corresponding 3D cone taper.

#### Three-dimensional simulation

##### Conservation of mass

Physically, mass diffusing in any domain should be constant, however the finite limits of computer precision and large numbers of voxels in 3D simulations allow the opportunity for round-off error to accumulate.

To quantify this effect for the serial (1 thread) simulation, we simulated diffusion on a Y-shaped geometry consisting of three sections, each of length 10 μm and diameter 2 μm. One section is positioned from (0, 0, 0) to (10, 0, 0). The other sections continue from there to 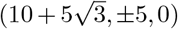, i.e. 30 degree changes in either direction. While the exact orientation would have no effect on 1D chemical or electrical simulation, we specify these details because they affect the exact number of voxels in the 3D simulation as narrow angles would lead to more overlap of the logical shapes in 3D space and hence less total voxels. We used the default discretization with voxels with dx=0.25 μm on each side, for a total of 7,904 voxels in our model. Substance diffused with diffusion constant of 1 μm^2^/ms) starting from a concentration of 1 μM on the section parallel to the *x*-axis and 100 nM on the other two sections and ran for varying lengths of time. For fixed step simulation, we started with a timestep of *dt* = 0.025 ms, NEURON’s default. By *t* = 100,000 ms, (i.e. after 4 million timesteps) conservation error accumulation led to a relative change of about 8.2219 × 10^−10^ of the total mass. As timestep reduced to *dt* = 0.0125 ms and *dt* = 0.00625 ms, the relative change in mass after 100,000 ms reduced to 7.004 × 10^−12^ and 1.7741 × 10^−13^, respectively. For variable step, using NEURON’s default tolerance and an `atolscale` of 10^−6^ for the state variable. Without scaling, NEURON’s default error tolerance would be 1 μM, small enough for sodium and potassium, but far too large for physiological concentrations of e.g. calcium which are often about 50-100 nM (Grienberger and Konnerth, 2012). With these settings, by *t* = 100,000 ms, variable step integration accrued a relative change of 2.7389 × 10^−13^ of total mass over 33,233,603 timesteps. We note that in practice, many NEURON simulations run for orders of magnitude less time, and can expect even smaller error accumulation.

We further examined conservation of mass by running the same simulation with 4 compute threads. The relative errors in both fixed step and variable step matched the results reported above for the serial case.

##### Ion channel fluxes

To examine the interplay between membrane potential, ion channels, and diffusion, we simulated sodium dynamics at various diffusion rates within a cylindrical “soma” geometry 10 μm in length and 10 μm in diameter with Hodgkin-Huxley channels under a continuous current injection of 0.1 nA. This current injection was sufficient to cause the cell to fire a train of action potentials, each of which admits sodium current into the cell, raising the sodium concentration. (Sodium concentration change is only simulated when sodium dynamics are explicitly modeled, either through NEURON’s rxd mechanism as here or through certain MOD file mechanisms.) As shown in Figure 5, with a low sodium diffusion rate, the sodium concentration near the surface builds up rapidly. As the diffusion rate increases, the surface concentration approaches that of the corresponding 1D simulation (not shown) as the sodium is more able to spread across the dendrite’s cross-section. By definition, the difference in sodium concentration in the surface voxels leads to a difference in the sodium Nernst potential (which is automatically recalculated by NEURON), which affects subsequent sodium currents and hence spike timing and shape, leading to the separation of spike times for the different diffusion rates shown in the inset to Figure 5A.

**Fig. 5.**
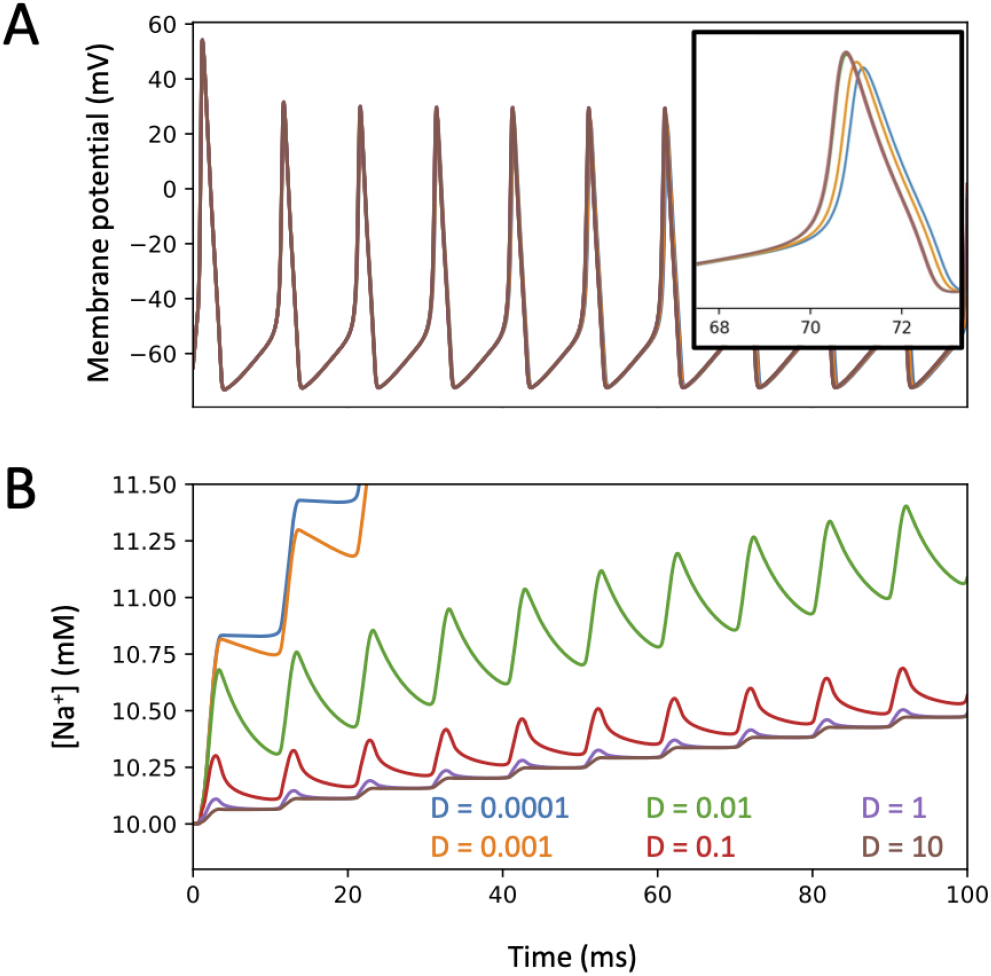
(A) Membrane potential and (B) surface voxel sodium concentration ofa 3D cylindrical soma with Hodgkin-Huxley channels and sodium accumulation, 10 μm in diameter and 10 μm in length with a constant current injection of 0.1 nA at various diffusion constants D (μm^2^/ms). Inset: small but visible separation of the action potential spike traces between the simulations with the two smallest diffusion constants and the rest by 70 ms into the simulation.

In the case of a single section, the same dynamics would be observed for a 2D model using radial shells to incorporate the difference between near-plasma-membrane concentrations and interior concentrations, however the 3D approach used here avoids the non-physical-realizability of radial shells at branch points (see e.g., Figure 1 in Chen and De Schutter (2017)).

###### 3D simulation on realistic geometry

For a more complete test, we compared simulations of scalar bistable dynamics on a realistic cell morphology using our algorithm with using the 3D cell biology simulator VCell (Cowan et al., 2012, Schaff et al., 1997). We used CTNG in NEURON to voxelize the morphology of NeuroMorpho.Org:NMO_02699 (Ascoli et al., 2007, Nikolenko et al., 2007). The voxelized data was exported to a stack of PNG images, where each image represents a *z*-slice with a value of 0 for voxels not in the cell and a value of 255 for voxels in the cell. These image stacks were then loaded into VCell with each pixel corresponding to one voxel. We note, however, that while this transfer approach correctly transfers information about which voxels are included, it loses the fractional volume calculated for surface voxels within NEURON, so the two tools are not expected to produce identical results as the boundaries vary slightly. In each tool, the initial concentrations were set to be 1 mM in the distal apical and 0 mM elsewhere. Reaction-diffusion was simulated until time *t* = 220 ms, and corresponding *z*-slices were compared. With both tools, the wavefront was at the same approximate location mid-soma and showed comparable curvature; Figure 6.

**Fig. 6.**
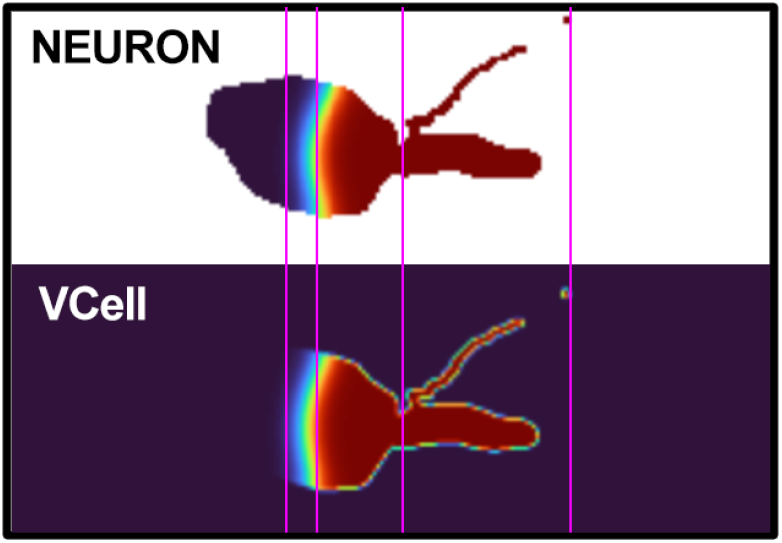
NEURON vs VCell Comparison. Reaction-diffusion NEURON and VCell simulation results of one cell z-slice at *t* = 220 ms. Image cropped to show relevant cell slice areas. Note the similarity between the characteristics of the wave curvature, approximate wave position, and the thickness ofthewave front in each simulation.

##### Orientation sensitivity with propagating wave

The orientation of a section affects how many voxels will be on the boundary and how the surface cuts through them, but the boundary voxel partial volumes and surface areas are inherently only approximations. To assess the impact of these approximations on models incorporating both 3D reaction and diffusion and components that are sensitive to 1D concentrations (e.g. ion channel kinetics specified using NMODL; Hines and Carnevale (2000)), we considered wave propagation governed by the the scalar bistable equation, *u_t_* = *D*Δ*u* – *u*(1 – *u*)(*α* – *u*), and timed wave propagation based on 1D concentrations. Here Δ is the Laplacian operator, *D* is the diffusion constant, and *α* is a threshold concentration, above which in the absense of diffusion concentrations will tend to increase and below which concentrations will tend to decrease. As observed in McDougal et al. (2013b), these dynamics exhibit key characteristics of some intracellular signaling processes, like calcium waves. Furthermore, this equation has a known analytic solution in the 1D infinite line case that can be used for validation.

To quantify this effect, we tested 100 random orientations of a 251 μm long dendrite of diameter 2 μm. We initialized the wave with a concentration of 1 on the first 50 μm and 0 elsewhere, then let it diffuse with a diffusion constant of *D* = 1 μm^2^/ms. (Concentration by default in NEURON is represented in mM, however the units are omitted here as the dynamics are the same as long as the units are consistent.) For each orientation, we estimated wave speed for three different choices of *α* (0.15, 0.25, 0.35) and three values of dx (2^−1^, 2^−1.5^, 2^−2^ μm). A plane wave in an infinite cylinder with these dynamics is known to propagate with a wave speed of 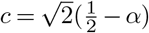 (see e.g. Fife (1979)). We estimated the wave speed in each simulation by measuring the time it took for the wave front (defined as the farthest point with an average 1D concentration over 0.5) to move from position 100 to 200 μm. These positions and the total length of the dendrite were chosen as they were found to allow reasonably accurate approximations of the wave speed in 1D simulations – i.e. a large enough distance to be free of boundary effects – while keeping the geometry small enough that the 900 total 3D simulations involved in this study could be run in a reasonable amount of time.

For *α* = 0.25, the average relative error in the estimated wave speed decreased proportionally to dx (4.59 ± 2.24 % for dx=2^−1^, 3.12 ± 1.56 % for dx=2^−1.5^, and 2.17 ± 1.09 % for dx=2^−2^). At NEURON’s default resolution of dx=0.25 μm, all orientations led to less than 4% relative error in estimated wave speed; about three-quarters (74 out of 100) showed less than 3% relative error, and about one-fifth (21 out of 100) had less than 1% relative error, with the minimum being 0.13%; Figure 7. All three values of *α* tested showed similar distributions of relative errors of wave speed (not shown). Cylinders whose axis was parallel to the *x, y* plane or mostly vertical gave less error in wave-time estimates than cylinders whose axes were not aligned with the Cartesian grid.

**Fig. 7.**
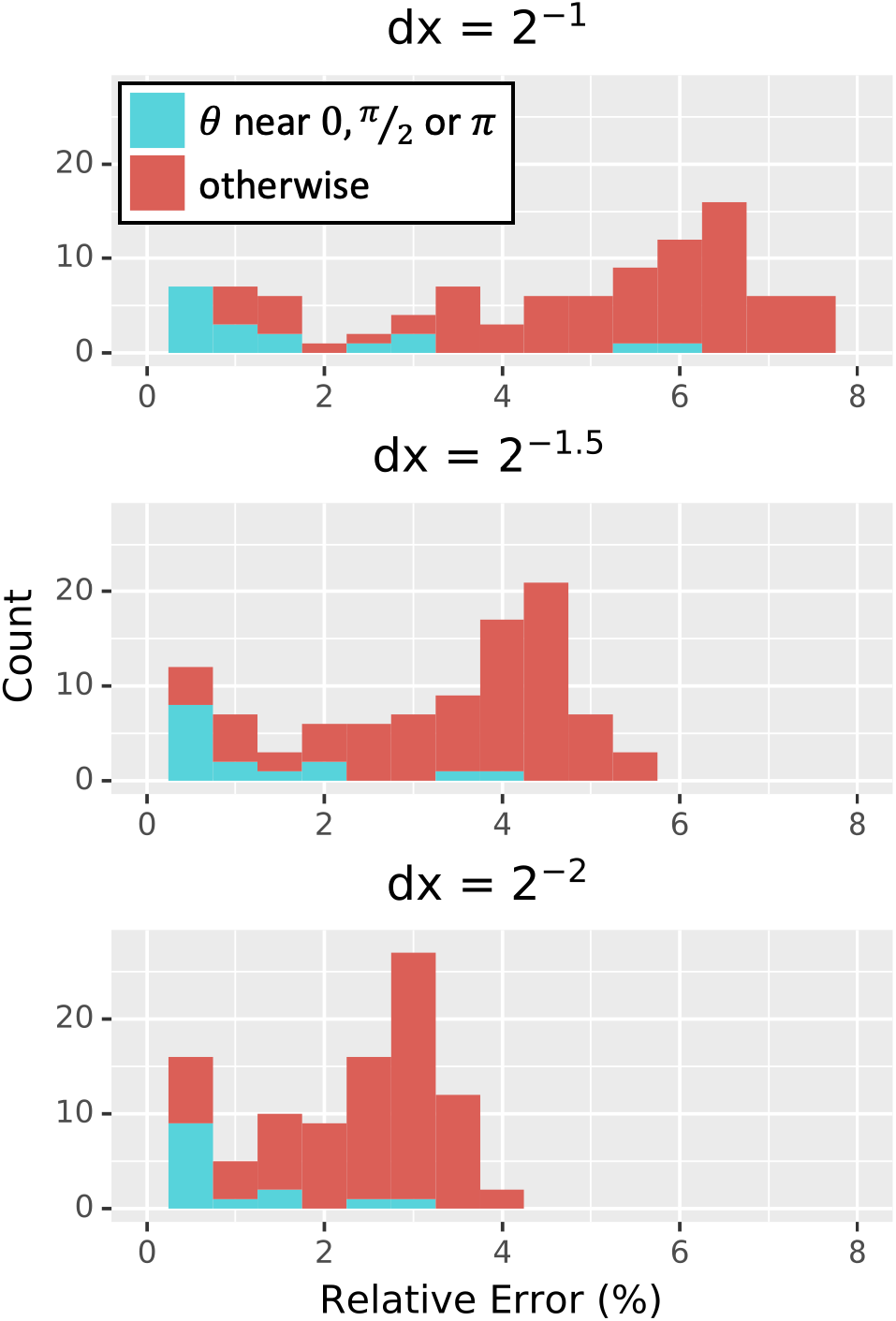
Distribution of relative error in wave speed for the scalar bistable wave with *α* = 0.25 derived from 100 random orientations of a 251 μm long, 2 μm diameter cylinder simulated in 3D at three choices of dx (measured in μm); all graphs are on the same scale. Simulations with *θ* within 0.1 radians of 0, *π*/2, or *π* corresponding to cylinders nearly parallel to either the *x, y* plane or the *z* axis are in cyan and tended to have lower errors than those that were oriented otherwise.

#### Hybrid 1D-3D simulation conservation of mass

Simulations of diffusion should conserve mass for 3D and hybrid 1D-3D models. To test hybrid conservation, simple hybrid models were used, where one section joined to one or two other sections, either aligned or a Y-shaped join. A region of initially elevated concentration was placed in one section away from the join and diffusion to the neighbouring sections was simulated. Using different voxel sizes and time-steps showed similar change in total concentrations, on the order of 10^−11^ % of initial amount, consistent with the expected numerical error.

### Performance

Defining and simulating a 3D model are logically separate activities: a model only needs to be defined once to be simulated many times (e.g. with different parameters). The most time-consuming part of the definition phase is the voxelization process. Furthermore, in principle, any voxelization that generates the appropriate data structures and maps voxels to segments could be used by the simulation engine. As such, we measure the performance of voxelization (currently single-threaded; described in the Voxelization Section) and the performance of simulation (multi-threaded; described in the 3D Simulation Section) separately. To assess the performance using realistic cell shapes, we tested 21 randomly selected morphologies (listed in the Random realistic neuron morphologies Section) with realistic diameters and 3D data from NeuroMorpho.Org (Ascoli et al., 2007).

#### Voxelization

To assess the voxelization performance, we loaded each of the 21 randomly chosen neuron morphologies one at a time and recorded the initialization time and estimated volume for many choices of spatial resolution dx, typically from 0.05 μm to 0.5 μm. Each timing was run in a separate process, as NEURON caches the results to avoid voxelizing the same cell more than once (i.e. subsequent model initializations skip the voxelization step). As shown in Figure 8A, the time required to voxelize/discretize the cell scaled nearly consistently at about 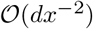 regardless of the morphology, with the main exceptions happening for large `dx`. As before, relative volume error was estimated using the volume calculated for the smallest measured `dx` as the estimated true volume. The relationship between estimated relative volume error as calculated in the convergence on a cylinder Section and time spent doing the discretization was noisy and less consistent across morphologies but generally scaled between 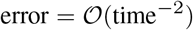 and 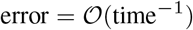 as shown in Figure 8B.

**Fig. 8.**
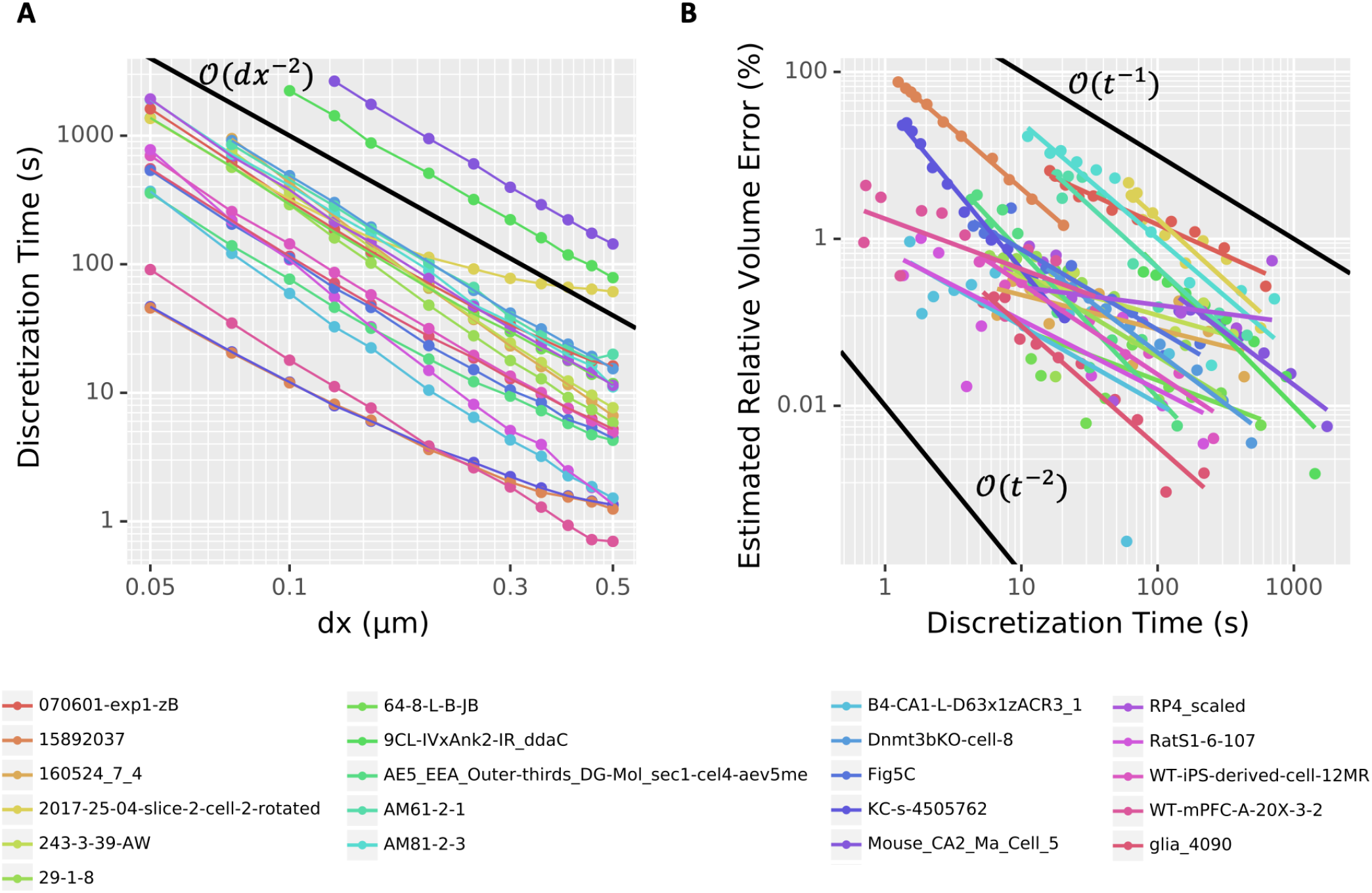
(A) Voxelization time scales as approximately 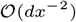. Dots denote measured data from 21 different morphologies from NeuroMorpho.Org; colored connecting line segments are illustrative only. (B) The scaling of estimated relative volume error varies depending on the morphology, but typically lies between 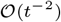 and 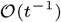, where *t* is the time spent computing the voxelization. Points denoted measured values; corresponding colored lines represent a linear (in log-log space) best fit. In both (A) and (B), black lines give examples of perfect scaling at the rate indicated.

#### 3D simulation

To assess the scaling of our simulation algorithm with the number of threads used, we simulated three different dynamics (pure diffusion, bistable wave, and calcium wave) across two morphologies (a cylinder of length 50 μm and diameter 1 μm and a reconstructed cell (NeuroMorpho.Org’s NMO_77436 (Canchi et al., 2017)), with three spatial resolutions (dx=0.25 μm, dx=0.125 μm, and dx=0.0625 μm). The pure diffusion dynamics were governed by Fick’s laws. The bistable wave modeled here implements the scalar bistable wave equation *u_t_* = *D*∇^2^*u* – *u*(1 – *u*)(*α* – *u*) analyzed in (Fife, 1979), and previously used as an example of reaction-diffusion phenomena in (McDougal et al., 2013b). The calcium wave model implements *Ca*^2+^-induced-*Ca*^2+^-release (CICR) driven by the endoplasmic reticulum, and is a simplified version of (Neymotin et al., 2015). Waves were initiated by an area of elevated cytosolic concentration (*u* for the bistable wave and IP3 for the calcium wave) in the first 25 μm in the cylinder case and in section `dend_7[19]` of the apical dendrite which starts approximately 9.35 μm from the soma in the morphologically detailed case.

Excluding the cylinder diffusion and cylinder bistable wave on the coarsest resolution (dx=0.25 μm), which both initially ran in under 1 second (and for whom threading overhead is thus non-trivial), the rest of the simulations showed speedup as the number of threads increased; Figure 9. For the 16 other combinations of morphology, model, and dx: using 4 threads reduced runtime by up to a factor of 3.63 (2.00 ± 0.65 on average); using 8 threads reduced runtime by up to a factor of 6.39 (3.20 ± 1.29 on average); using 16 threads reduced runtime by a factor of up to 9.76 (5.07 ± 2.28 on average). The reported runtimes here are based on the best of three runs to limit the contribution from background tasks.

**Fig. 9.**
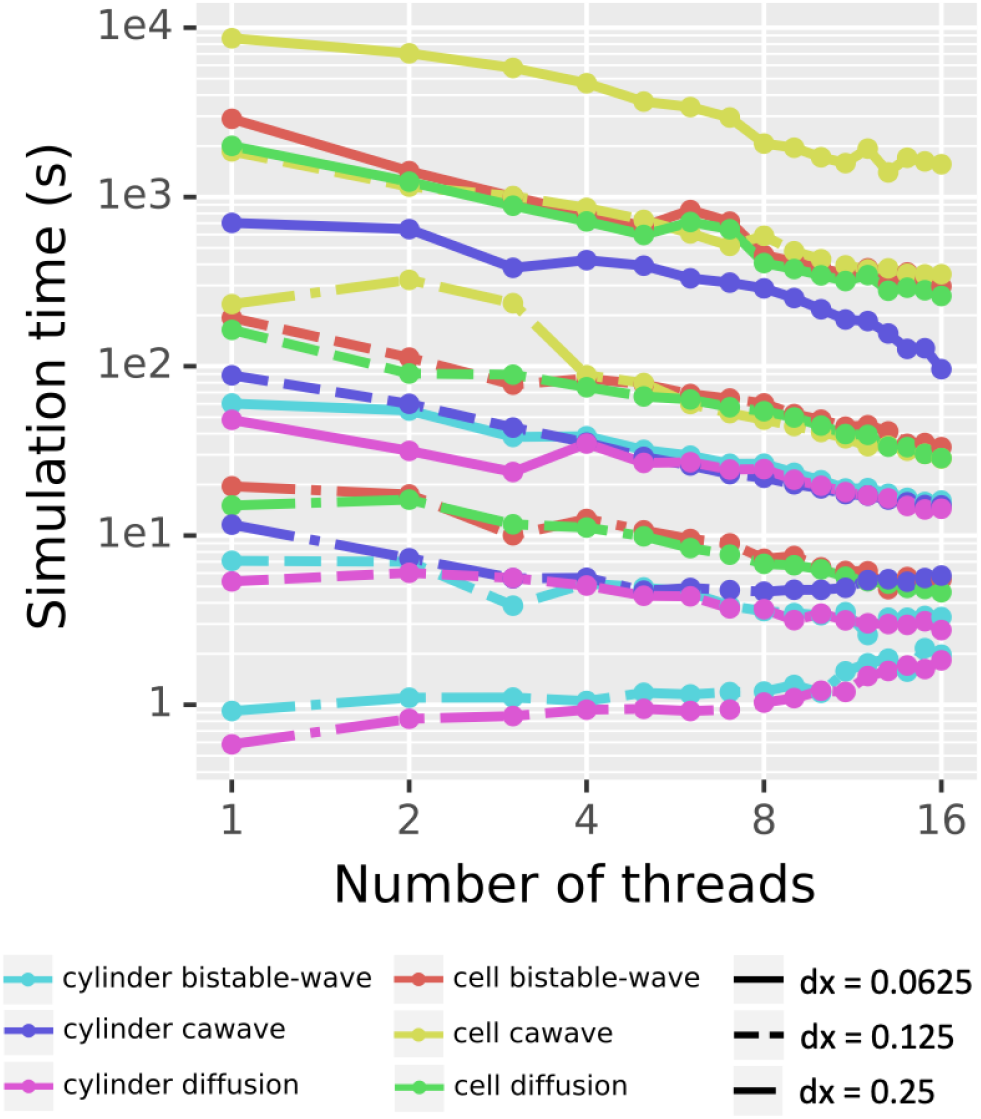
Parallel scaling. Testing all 18 combinations of 2 choices of morphology, 3 choices of model, and 3 choices of dx shows decreased real time for simulation for most combinations as the number of threads increases from 1 to 16. Pure diffusion and a bistable wave on a cylinder at dx=0.25 μm were the only two test cases that took more real time with 16 threads than with 1 thread. Solid lines indicate dx=0.0625, dashed lines dx=0.125, and dash-dot lines dx=0.25.

Reported times exclude voxelization time, analyzed above, which is currently single-threaded and is only performed once for a given morphology regardless of the number of simulation studies performed.

To further test performance speed-up, we experimented with different settings of cache prefetch, which in our experiments ultimately did not show significant difference in parallel scaling.

### Examples

Three-dimensional simulation offers the ability to explore both the role of a neuron’s three-dimensional shape – which is especially relevant wherever the neuron is not approximately conical, such as where the dendrites meet the soma or a spine connects to a dendrite – and the role of precise spatial positioning for e.g. synapses. In this subsection, we examine examples of each of these, discuss relevant implementation details, and examine different visualization strategies for the resulting volumetric data.

#### Dendrite-soma intersection

Certain cellular phenomena are typically found in one region of the cell and typically not present in neighboring regions even when the neighboring regions are known to be able to support the phenomena. For example, waves of elevated intracellular calcium in pyramidal cells observed in apical dendrites only sometimes invade the soma but when they do are capable of propagating across the soma (Hagenston et al., 2008). There exist mathematical models of wave phenomena where it is known that domain geometry affects wave propagation (e.g. Dronne et al. (2009)); 3D simulation allows us to study if the morphology plays a similar role in problems of neuroscientific importance.

For example, we simulated the scalar bistable equation with a threshold *α* = 0.1 mM in the morphology of Neuro-Morpho.Org:NMO_53113 (Ascoli et al., 2007, Malik et al., 2016) starting with a concentration of 1 mM on the distal apical and 0 mM elsewhere, and a diffusion constant of 0.25 μm^2^/ms. No other dynamics were included; in particular, no ion channels were simulated and there was no flux across the plasma membrane (Neumann boundary conditions). We chose this cell in part because the soma of this cell was specified using a soma outline in ASC format, allowing NEURON to construct a non-cylindrical approximation to the soma shape. Using a fixed-step simulation (dt=0.25 ms), we simulated the volume containing the soma and all sections whose center was within a path distance of 70 μm from the soma’s center in 3D, with the rest of the cell in 1D. Within this volume of 3D simulation, the smallest diameter was 0.18 μm, and we used a dx = 0.17 μm. Under these conditions, a wave of elevated concentration propagated from the apical toward the soma at approximately uniform speed. Near the soma, the wave front curved and slowed, but propagated into the soma where it eventually straightened and resumed its initial speed; Figure 1A shows the progression of the wave front over time using contours on a 2D projection of the cell.

To assess if the hybrid approach was accurately simulating wave behavior within our region of interest near the soma, we repeated the experiment using all sections whose center was within a path distance of 100 μm from the center of the soma; this expansion added an additional 19 sections to the 3D domain. Simulating in 3D on this expanded domain gave a visually identical contour map of wave propagation (not shown), and an identical prediction for when the wave would cross the center of the soma (t=188.255 ms), defined as the first time the 1D concentration at the center of the soma exceeded the half-maximal value. This consistency suggests that our original simulation was not losing significant accuracy near the soma despite simulating distal parts of the cell in 1D. By contrast, simulating only the soma and the sections directly connected to the soma in 3D led to a different wave crossing time (t=194.58 ms), which therefore indicates this smaller region is not a sufficiently large 3D region for studying behavior near the soma.

We note that Figure 6 presents a similar experiment on a different morphology showing a colorcoded 2D slice at a specific time point. The visualization in the latter figure shows more detail on the concentration distribution near the wave front, but cannot show the propagation of the wave front over time.

From this example, we make a few observations that apply to any such experiment: (1) areas that are far from the region of interest do not need to be simulated in 3D; when studying effects at the dendrite-soma intersection, this allows larger dx values than would be possible if the fine distal dendrites also needed to be simulated in 3D. (2) Conversely, such experiments are fundamentally about the role of boundary conditions, therefore other boundaries must be at a far enough distance from the region of interest so as not to affect the results. (3) The accuracy of any results arising from such simulations depends on the accuracy of the voxelized reconstructions. For this reason, we currently recommend using reconstructions in ASC format with a soma outline, as this will provide a non-cylindrical soma. Future versions of NEURON will allow importing a predefined voxelization to more accurately reflect the observed shape, but this will necessarily require matching the surface areas and volumes on the 3D chemical domain with that used for electrophysiology simulation. (4) Regenerative waves have a leading edge that can be plotted on a contour map at regular intervals showing the progression of the wave over time.

#### Spines

Numerous publications have examined the interactions of spines with each other (e.g. Chiu et al. (2013)), how concentrations may be compartmentalized within spines (e.g. Yuste et al. (2000)) and the relationship between membrane potential in spine and dendrite (e.g. Jayant et al. (2017)). As with dendrites meeting the soma, the exact nature of dynamics at the spine-dendrite juncture depends on the shape of this connection. This is in part dependent on the angle with which the spine attaches to its dendrite, a detail completely lost in 1D simulation, but analagous to the issues arising at the soma.

Spine-dendrite modeling, however, also introduces a new challenge arising from non-physically realizable models where the same volumes is part of two separate sections. Such overlapping sections are common in NEURON, as sections are by default connected to the centroid of the parent section. For most models, the length of most sections is longer than the length of the diameter and the child diameters are generally comparable to the parent diameters, so any discrepancies in local surface area or volume due to the overlaps are typically minimal. Models with spines are a notable exception; spine necks vary in shape and size but for example in layer 6 pyramidal cells of the mouse somatosensory cortex are typically less than 0.2 μm in diameter and less than 2 μm long (Ofer et al., 2021), and so attaching the spine at the centroid places much of the neck inside the dendrite. Here our choice of mapping the voxel to the 1D compartment closest to the presumptive soma assures that the spine neck is only that portion extending beyond the dendrite proper, but the 3D volume and surface area calculations will be based only on the part that extends beyond the dendrite, and thus the volumes will disagree, and it is possible that some segments may not have any true surface area. This discrepancy between the 1D and 3D representations can be mitigated by shifting the start of the spine neck to be some distance (almost a radius) away from the centroid of the parent dendrite using the appropriate NEURON `pt3dstyle` while keeping the perimeter inside the parent dendrite. In the case of a cylindrical dendrite with a smaller orthogonal spine, the maximum spine distance from the dendrite centroid can be found by considering the circular cross-section of the dendrite meeting the rectangular crosssection of the spine neck, placed inside the dendrite such that the two lower vertices are on the perimeter of the dendrite. The resulting distance *d* from the centroid is determined by a right triangle, formed by the center of the dendrite, one of the lower vertices of the spine neck, and the center of the lower edge of the spine neck. This gives a triangle with hypotenuse *r_d_*, adjacent *r_n_* and opposite *r_d_ – d*, where *r_d_* is the radius of the dendrite and *r_n_* the radius of the spine neck. Then by Pythagoras’s theorem 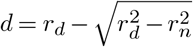.

Using this rule, we constructed a cylindrical dendrite 2.5 μm in diameter and 6 μm long. We attached two spines at position 3 μm orthogonal to the dendrite with necks of length 3 μm, diameter 0.1 μm and cylindrical heads of length 0.5 μm and diameter 0.6 μm. To simulate the spread of a substance from the spines, they were initially filled with substance to a concentration of 2 mM, with 0 mM in the dendrite. When the substance was allowed to diffuse at a rate of 0.01 μm^2^/ms, the dynamics of the concentrations within the dendrite varied depending on the angle separating the two spines; Figure 1B. We considered two cases: spines 30 degrees apart, and spines 180 degrees apart. We note that in a non-3D simulation these two cases would give identical results. The peak dendritic concentration in each case was reached within the first 0.5 ms, with the closer spines leading to a peak concentration 89% as high as with the spines on the opposite side of the dendrite. All voxels dropped below a concentration of 0.15 mM 17.575 ms earlier when the dendrites were near each other than when they were opposite each other. If the threshold for triggering another reaction was around 0.15 mM, this difference in time above that value could make the difference between whether or not the downstream reaction was triggered. For different choices of parameters (e.g. with a thinner dendrite), the same model could have the peak concentration drop below the threshold in the other order.

#### Three-dimensional localization of synapses

Metabotropic receptors and other mechanisms exist at specific points in 3D space. Such dynamics in 1D are often specified using files written in NMODL (Hines and Carnevale, 2000), a domain specific language for ion channel, receptor, and artificial cell kinetics supported by NEURON, Arbor (Akar et al., 2019), and the Python nmodl module (github.com/bluebrain/nmodl). We can apply the same approach to synapses located in 3D space, but a few extra considerations are necessary.

First, we begin by defining our post-synaptic response kinetics. In principle, these can be arbitrarily complicated to reproduce experimental observations, however as a first approximation it is not uncommon in modeling to see mechanisms where the rate of production jumps abruptly in response to synaptic and decays exponentially; such an NMODL file is shown in Fig. 10. In this file, `g` denotes the rate of production of a substance; physically, this corresponds to a change in mass per ms. To support traditional NMODL files, all currents generated by an NMODL mechanism are distributed over the entire segment surface. As a segment is the smallest electrical compartment, that behavior is correct for the electrical aspects of the simulation, however distributing e.g. sodium currents across the surface would result in sodium changes in all surface voxels. To avoid this issue, a 3D targeted NMODL mechanism must generate only `NONSPECIFIC_CURRENT` with chemical changes driven solely by the rate `g`.

**Fig. 10.**
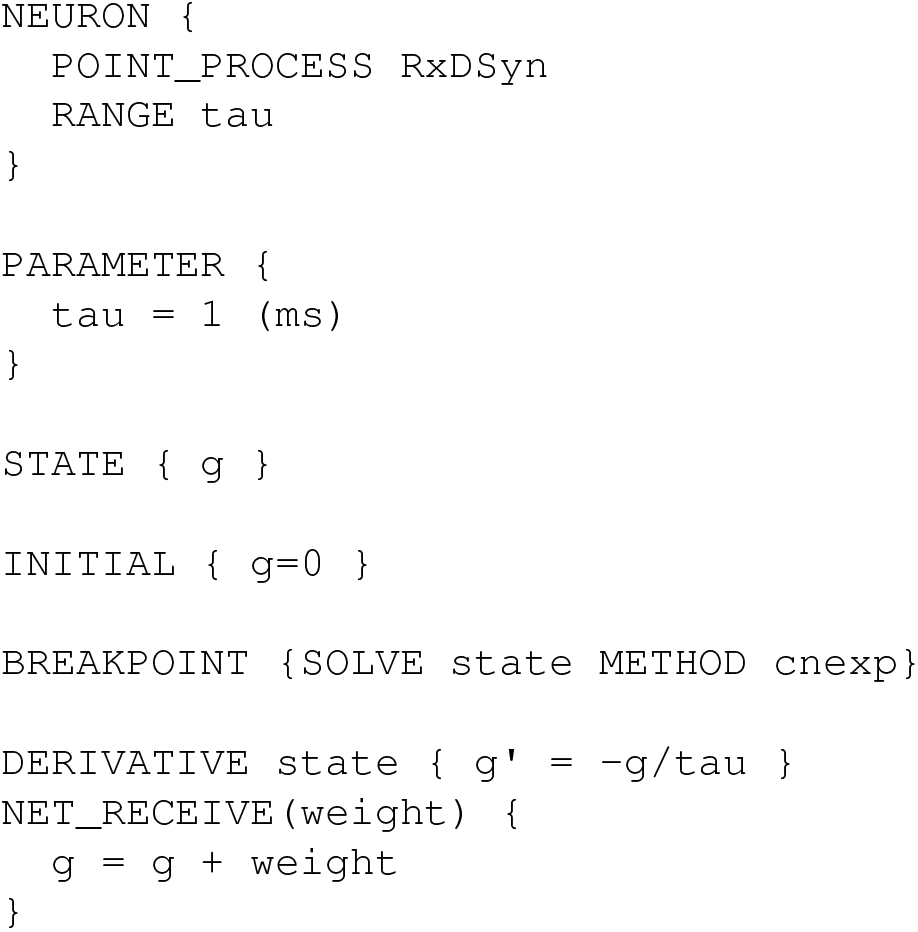
Source code for a generic NMODL mechanism called `RxDSyn` that receives synaptic events (`NET_RECEIVE` block) causing the flux *g* to increase abruptly in response to an event by an associated weight, with the flux decaying exponentially with time constant tau thereafter (`DERIVATIVE` block).

NMODL mechanisms must be compiled before they can be used. This is typically done by running `nrnivmodl`, but we note additional compilation options are sometimes available. NEURON loads compiled NMODL mechanisms from the current directory at startup and can also load them on demand via `h.nrn_load_dll`. Once loaded, the `POINT_PROCESS` name (`RxDSyn` in Fig. 10) is available as a class in NEURON’s `h` object. That is, a new instance could be created by `r = h.RxDSyn(seg)`, where `seg` is the segment that contains the mechanism.

Once we have picked the kinetics, the next step is to identify the 3D location to place them. If `ca` is an `rxd.Species` on a 3D region, then `ca.nodes((x, y, z))` is an `rxd.NodeList` of `ca` nodes containing the point `(x, y, z)`. As each node covers a volume, there are many points within a `Node` but at most one `Node` that contains the point unless the `rxd.Species` is present on more than one region (e.g. calcium might be present in both the ER and the cytosol, as in Neymotin et al. (2015)). The coordinates of the center of a `Node`’s voxel are (`node.x3d, node.y3d, node.z3d`). Note that if `node` is on the surface, then `node.surface_area` should be strictly positive. If the surface exactly touches a grid corner, it is possible that some voxels with zero surface area will be included in the mesh, but as such, these should not be used for surface-based kinetics. If the segment containing the mechanism was initially unknown, it can be obtained from the selected node via `node.segment`.

Mechanisms may be connected to one or more nodes by passing a pointer to the rate to the node’s include_flux method; in our example, this is `node.include_flux(r._ref_g)`. By default, this method assumes g is measured in molecules per second; these units ensure that the same total amount of substance is enters the cell regardless of the discretization. The same flux rate may optionally be applied to other nodes.

Once this is done and the post-synaptic dynamics are connected to a presynaptic event source (e.g. a membrane potential crossing a threshold or a random spike train), then presynaptic events will trigger production of the `node`’s substance at a rate that decays over time (if using the kinetics of Figure 10) and that substance is free to diffuse away; Figure 11.

**Fig. 11.**
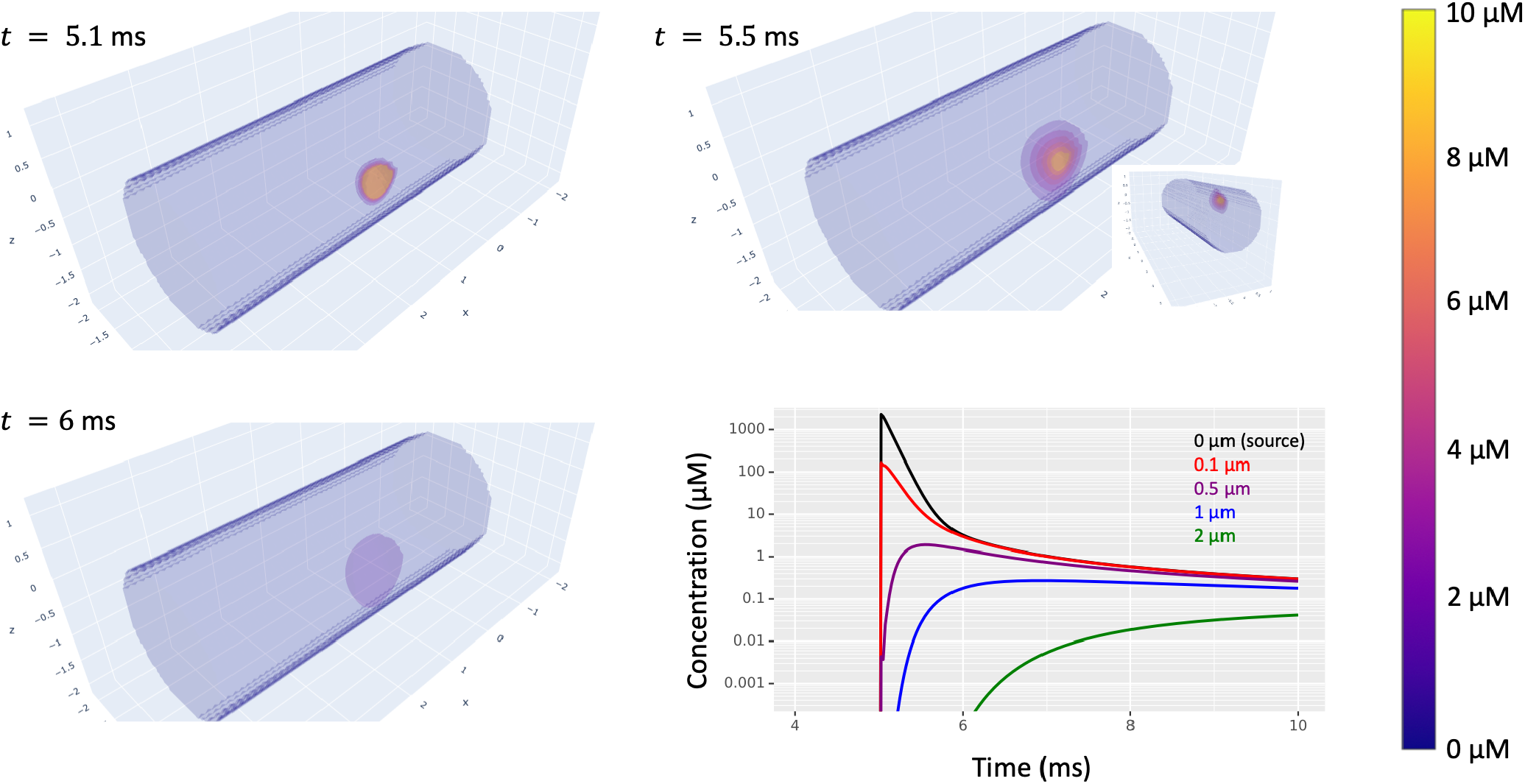
Simulated diffusion in a cylindrical dendrite of a substance produced at (2.5, 0.975, 0.275) in response to synaptic input at time *t* = 5 ms. The volumetric images highlight translucent level sets with all concentrations above 10 μm displayed the same. Bottom-right: concentration (on a log scale) vs time at 5 distances from the source on the surface of the dendrite. Inset: a volumetric view from a different angle showing the extent of diffusion into the interior of the dendrite.

## Discussion

The development version of NEURON provides built-in support for parallel, 3D deterministic simulation of intracellular reaction-diffusion dynamics (e.g. protein and ion interactions and diffusion) in whole neurons and in modeler-selected Sections of interest; the remaining Sections, with kinetics expressed identically, continue to use 1D reaction-diffusion simulation, allowing computational resources to be targeted toward locations where the 3D shape is likely to matter such as the relatively large volumes near the soma. Selected cells or Sections are voxelized using an updated version of the CTNG algorithm (McDougal et al., 2013a) that exploits convexity. Synapses can optionally target their effects to specific 3D compartments (e.g. production of a certain mass of messenger from activation of a metabotropic synapse). Electrophysiology simulations remain simulated as branching 1-dimensional sections as is appropriate given their larger space constants.

### Alternative strategies

There are two main alternative approaches in the literature for combining 3D reaction-diffusion kinetics with electrophysiology.

The first alternative approach is to have an integrated solver that uses a single mesh. STEPS, for example, simulates ion channel, etc. activity on the surface of the 3D mesh (Hepburn et al., 2013); a similar approach was used in the Virtual NEURON study (Brown et al., 2011). Using the same mesh eliminates the possibility of numerical artifacts from coupling, automatically ensures consistent surface areas, and eliminates the possibility of interior surface (e.g., from spines mis-connected at the centroid). We have avoided this approach, instead using 1D electrical with 3D reaction-diffusion as in (Grein et al., 2014) to allow the electrical dynamics to be consistent regardless of the dimensionality of the reaction-diffusion simulation, to take advantage of the 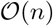 implicit simulation of electrical dynamics on such a 1D-structure (Hines, 1984), and for compatibility with the over 2000 existing NEURON models (i.e. extending an existing NEURON model with 3D intracellular reaction-diffusion dynamics does not require modifying the existing components, unless a change is desired to their behavior).

The second alternative approach is to use multisimulation; that is, to combine a solver specializing in ion channels and the cable equation like NEURON or MOOSE (Dudani et al., 2009) with an external solver specializing in reaction-diffusion simulation. We and our colleagues have used this approach for stochastic 3D model simulation with NEURON Time Warp (Lin et al., 2017). KappaNEURON likewise combines NEURON with the rule-based reaction-diffusion simulator SpatialKappa (Sterratt et al., 2014). Grein and colleagues used a similar approach for deterministic 3D simulation coupling NEURON with uG (Grein et al., 2014). Additionally, we note that NEURON supports the general multisimulation framework MUSIC (Djurfeldt et al., 2010) which has been used to connect MOOSE and NeuroRD (Brandi et al., 2011). The multisimulation approach is appealing as it allows each simulator to specialize in its own problem domain, providing a rich set of simulatable features, with each simulator programmed independently. While recognizing the benefits and flexibility of multisimulation, we chose to build our 3D intracellular simulation capability within NEURON to allow for a unified Jacobian matrix, allowing for variable step simulation, to provide consistent coupling semantics, to avoid the need for syncing data between two different potentially parallel tools, and to avoid the need for users to learn two simulator tools.

### Special considerations

Models with exactly zero diffusion of a species that enters or leaves through ion channels pose specific issues in comparing 1D and 3D simulations or 3D simulations with different discretizations. Although mathematically convenient, these models are non-physical as diffusion is necessary to bring a molecule to or through an ion channel. Concentration changes in 1D are based on the active geometry, typically the whole dendrite. Increasing the spatial resolution (e.g. tripling nseg) in a 1D model with no diffusion has no direct effect on concentration dynamics as the volume and total current both scale by the same fraction. In contrast, in a 3D model as ions must enter via the surface, with no diffusion, they are trapped there. Reducing voxel edge size by a factor of 2 changes the volume by a factor of 8 but the surface area contained in the voxel by only a factor of 4 on average, leading to a factor of 2 change in the rate at which concentration in surface voxels changes, which could affect ion channel kinetics.

In principle, multigriding could be extended to be used within the chemical dynamics in 1D or 3D; i.e. a species prone to steeper gradients could be simulated on a finer grid, but this risks introducing artifacts, especially in the case of slow or zero diffusion. For example, suppose molecule A on a coarse grid bound with molecule B from a refinement of the grid to form molecule AB. If AB is represented on the same fine grid, then when it dissociates A and B can return to their correct points of origin. If on the other hand, AB is represented on the coarse grid then when it dissociates B could end up in any of the corresponding fine meshes. Thus, even if the diffusion rate was set to zero for all species, molecule B could move by binding to A, entering the coarse grid, and then returning to a different fine grid compartment. A similar problem exists regardless of the relative sizes of the grids if they do not align perfectly. Meshless simulators, like MCell (Stiles et al., 1998), avoid this class of problems entirely at the cost of having to simulate each molecule separately. We note that this problem only pertains to overlapping meshes; separate mesh resolutions on different parts of the cell (e.g. large near the soma, smaller in the distal dendrites) are potentially compatible, although the mesh transition would not in general be expected to align to the boundary between Sections.

We note that the insight gained by a 3D simulation depends on the quality of the 3D mesh. CTNG or any of the alternative rules for converting point-diameter representations into a 3D mesh are inherently approximations as the full shape of the cell is under-determined by the reconstruction data. More importantly, however, we recommend that – regardless of metadata annotations – morphologies should be manually reviewed for slice artifacts (e.g. when we randomly selected 21 morphologies, we found that while none were strictly planar several showed minimal *z*-axis variation), for realistic and non-uniform diameters, for electrical connectivity (no pinch points where the diameter gets very small), and for *z*-axis errors (some reconstructions show abrupt changes in *z* values). There is no value in doing a 3D simulation if the 3D morphology is unrealistic.

3D time-series data is in general large and hard to visualize. We deal with the large volume of data by only automatically keeping the current state in memory. The time series of the concentration of a specific species at a specific compartment may be recorded using a `Vector`. For modelers needing to store or visualize all the states at a specific time, simulations may be stopped at a specific time point, where the states are then captured to an appropriate Python data structure. Throughout this paper, we have deliberately illustrated several approaches to visualizing such data: (1) line plots of a single species at a single point as in Figure 1B; (2) for traveling waves, plots of the location of the wave front at evenly spaced time points on a 2D projection or slice as in Figure 1A; (3) plots of the concentrations at the surface, analagous to the surface segment identities in Figure 4A; (4) heatmaps of a slice or projection at a specific time point as in Figure 6; and (5) translucent contour maps of concentration level sets at given time points as in Figure 11. Example Python code for each type of graph is available in this paper’s entry on ModelDB.

### Future directions and conclusions

NEURON is under continuous development. We intend to improve its support for 3D simulation by streamlining mesh generation: the CTNG algorithm is in-principle embarrassingly parallel and meshes and the data on them could in principle be saved and reused when relaunching NEURON. The first will require reimplementation of CTNG in pure C++ to avoid parallel limitations from Python’s GIL, and the second will require an efficient way of validating that the mesh aligns with the 1D skeleton. Additionally, we intend to integrate support for stochastic simulation to study more classes of dynamics and to more faithfully capture phenomena arising from very low concentrations or very small regions.

We believe that the approach described in this paper provides an intuitive way of incorporating intracellular reaction-diffusion dynamics in computational neuroscience models in a way that more faithfully captures the effects of geometry than is possible in a 1D or 1D + radial simulation. We hope that this allows new insights into the multi-scale processes that underlie our neural activity.

## Author contributions

CC and RAM designed the 3D simulation strategy in consultation with AJHN; CC implemented it in consultation with AJHN. LE and RAM designed the 3D voxelization strategy; LE implemented it. AJHN designed and implemented the surface subvoxelization, and improved simulation and voxelization robustness. RAM, HG, AJHN, and CC performed the analyses. All authors drafted, reviewed, and edited the manuscript.

## Acknowledgments

We thank Michael L. Hines and William W. Lytton for valuable discussions affecting the design of the interface and its implementation.

We thank the Yale Center for Research Computing for guidance and use of the research computing infrastructure, specifically the Farnam HPC.

